# Amplification and Suppression of Distinct Brain-wide Activity Patterns by Catecholamines

**DOI:** 10.1101/270645

**Authors:** RL van den Brink, S Nieuwenhuis, TH Donner

## Abstract

The widely projecting catecholaminergic (norepinephrine and dopamine) neurotransmitter systems profoundly shape the state of neuronal networks in the forebrain. Current models posit that the effects of catecholaminergic modulation on network dynamics are homogenous across the brain. However, the brain is equipped with a variety of catecholamine receptors with distinct functional effects and heterogeneous density across brain regions. Consequently, catecholaminergic effects on brain-wide network dynamics might be more spatially specific than assumed. We tested this idea through the analysis of functional magnetic resonance imaging (fMRI) measurements performed in humans (19 females, 5 males) at ‘rest’ under pharmacological (atomoxetine-induced) elevation of catecholamine levels. We used a linear decomposition technique to identify spatial patterns of correlated fMRI signal fluctuations that were either increased or decreased by atomoxetine. This yielded two distinct spatial patterns, each expressing reliable and specific drug effects. The spatial structure of both fluctuation patterns resembled the spatial distribution of the expression of catecholamine receptor genes: α_1_ norepinephrine receptors (for the fluctuation pattern: placebo > atomoxetine), ‘D2-like’ dopamine receptors (pattern: atomoxetine > placebo), and β norepinephrine receptors (for both patterns, with correlations of opposite sign). We conclude that catecholaminergic effects on the forebrain are spatially more structured than traditionally assumed and at least in part explained by the heterogeneous distribution of various catecholamine receptors. Our findings link catecholaminergic effects on large-scale brain networks to low-level characteristics of the underlying neurotransmitter systems. They also provide key constraints for the development of realistic models of neuromodulatory effects on large-scale brain network dynamics.

**SIGNIFICANCE STATEMENT:** The catecholamines norepinephrine and dopamine are an important class of modulatory neurotransmitters. Because of the widespread and diffuse release of these neuromodulators, it has commonly been assumed that their effects on neural interactions are homogenous across the brain. Here, we present results from the human brain that challenge this view. We pharmacologically increased catecholamine levels and imaged the effects on the spontaneous covariations between brain-wide fMRI signals at ‘rest’. We identified two distinct spatial patterns of covariations: one that was amplified and another that was suppressed by catecholamines. Each pattern was associated with the heterogeneous spatial distribution of the expression of distinct catecholamine receptor genes. Our results provide novel insights into the catecholaminergic modulation of large-scale human brain dynamics.

## INTRODUCTION

Neuromodulators are important regulators of physiological arousal and profoundly shape the state of neuronal networks in the cerebral cortex. Catecholamines, an important class of neuromodulators including norepinephrine (NE) and dopamine (DA), amplify the gain of neuronal responses to sensory input (Berridge and Waterhouse, 2003; Winterer and Weinberger, 2004; Jacob et al., 2013; Polack et al., 2013). Current models of catecholaminergic modulation posit that this increase in response gain amplifies the signal-to-noise ratio of sensory responses at the network level (Servan-Schreiber et al., 1990; Aston-Jones and Cohen, 2005; Eckhoff et al., 2009; Shine et al., 2018). An assumption common to these models is that catecholamines boost neural gain homogenously across the entire brain. This assumption is grounded in the widespread projections of the brainstem structures releasing these neuromodulators, in particular the locus coeruleus (LC), the main source of NE in the forebrain (Aston-Jones and Cohen, 2005).

However, other findings cast doubt on this assumption. First, there exists a multitude of different catecholamine receptors, and each has a distinct and heterogeneous distribution across cortical areas (Zilles and Amunts, 2009; Nahimi et al., 2015). And second, several of these receptor types exhibit distinct functional effects on cortical state (McCormick et al., 1991; Ramos and Arnsten, 2007; Robbins and Arnsten, 2009; Noudoost and Moore, 2011; Salgado et al., 2016). As a consequence, the effects of catecholamines on neural dynamics might be more spatially specific than traditionally assumed, perhaps even with opposing signs between different sets of brain regions. Here, we tested this idea by imaging the spatial distribution of catecholamine-induced changes in large-scale human brain dynamics, and relating the resulting patterns of brain dynamics to the spatial distribution of several catecholamine receptor types.

fMRI signals fluctuate strongly in the absence of changes in sensory input and motor output (often called ‘resting-state’), and these fluctuations correlate between distributed brain regions (Biswal et al., 1995; Fox and Raichle, 2007; Schölvinck et al., 2010). In the following, we refer to this phenomenon as intrinsic fMRI signal correlations, or simply, correlations. We have previously examined the effect of increasing central catecholamine levels on intrinsic fMRI signal correlations in a double-blind placebo-controlled crossover design using the a NE transporter blocker atomoxetine (van den Brink et al., 2016). Atomoxetine increases central NE and DA levels (Bymaster et al., 2002; Devoto et al., 2004; Swanson et al., 2006; Koda et al., 2010). This revealed reductions in the strength of correlations across several spatial scales of brain organization: in summary measures of brain-wide coupling derived using graph theory; coupling between large-scale functional ‘networks’ as defined in resting-state fMRI studies (Fox and Raichle, 2007); and in a select set of brain regions in the occipital lobe. This reduction was a surprising effect. However, our previous analyses also had two important limitations. First, our previous study revealed only the prevailing catecholamine-induced changes in correlations and thus left open the possibility that atomoxetine, in addition to decreases, also induced weaker, or less widespread, increases in correlations. Second, due to the use of a pre-defined network parcellation scheme and summary statistics, our previous study could not uncover more fine-grained spatial patterns of spontaneous signal fluctuations that were amplified or suppressed by catecholamines.

Here, we re-analyzed our dataset (van den Brink et al., 2016) with a previously validated analysis approach (Donner et al., 2013) that was tailored to address both issues above. Our new analysis enabled us to (i) assess the spatial specificity and fine-grained neuroanatomical structure of cathecolaminergic modulation patterns; and (ii) quantify their spatial correspondence with the distribution of the expression of catecholamine receptor genes, as revealed by a unique brain-wide transcriptome database (Hawrylycz et al., 2012; Hawrylycz et al., 2015). The analysis identified two spatial patterns of fMRI signal correlations that were most strongly affected by catecholamines; one with increased correlation strength and the other with reduced strength. These distinct networks were each associated with the expression pattern of distinct catecholamine receptors.

## MATERIALS AND METHODS

### Participants and experimental design

We reanalyzed data from van den Brink et al. (2016). This dataset comprised eyes open ‘resting-state’ (blank fixation) fMRI measurements in 24 healthy human participants (19 females, 5 males). In each of two separate sessions, scheduled one week apart, two fMRI measurements were performed, one before and one after intake of either placebo or atomoxetine (40 mg). The study had a double-blind placebo-controlled crossover design, and was approved by the Leiden University Medical Ethics Committee. All participants gave written informed consent before the experiment, in accordance with the declaration of Helsinki. Salivary markers of central catecholamine levels confirmed drug uptake (Warren et al., 2017).

### MRI preprocessing

A full description of scan parameters and preprocessing details can be found in van den Brink et al. (2016). In brief, we applied the following preprocessing steps to the fMRI data (TR = 2.2 s; voxel size = 2.75 mm isotropic): realignment and motion correction; B0 unwarping; high-pass filtering at 100 s; prewhitening; smoothing at 5 mm FWHM; coregistration of the functional scans with an anatomical T1 scan to 2 mm isotropic MNI space; artifact removal using FMRIB’s ICA-based X-noiseifier (Griffanti et al., 2014; Salimi-Khorshidi et al., 2014). We recorded heart rate using a pulse oximeter and breath rate using a pneumatic belt during data acquisition. Between-condition differences in heart rate and breath rate were examined using *t*-tests. We applied retrospective image correction to account for differences in heart and breath rate between the atomoxetine and placebo conditions (Glover et al., 2000). In the current article, we primarily focus on the runs following atomoxetine / placebo ingestion, but use the pre-pill conditions as a baseline in control analyses.

### Brain parcellation

We extracted the fMRI time series of individual brain regions using the Automated Anatomical Labeling (AAL; Tzourio-Mazoyer et al., 2002) atlas, which contained 90 regions (cf. van den Brink et al., 2016). In control analyses, we also used a more fine-grained atlas that was based on a functional parcellation (Craddock et al., 2012). This atlas contained 140 individual brain regions. Overall, the Craddock atlas yielded highly similar results as the AAL atlas, in terms of both the direction and significance of effects. Thus, our primary analyses are based on the AAL atlas, while our findings with the Craddock atlas are reported as a control analysis at the end of the Results section.

### Inter-regional covariance of fMRI signal fluctuations

After averaging across voxels within each atlas-level brain region, we Z-scored the multivariate time series (*M*, with dimensionality imaging volumes by brain regions) for each run *i* and then computed the group-averaged covariance matrices (C) for the placebo and atomoxetine conditions (subscript P and A, respectively) via the following:

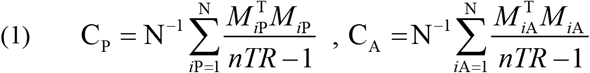

where *nTR* was the number of volumes (211), N was the number of participants (24), and ^T^ denoted a matrix transposition. The matrices C_P_ and C_A_ represented the covariance between the BOLD time series of all brain regions, averaged across participants. Note that by Z-scoring the time series, the units of C (covariance) are equivalent to the Pearson correlation coefficient.

### Eigenvalue decomposition of covariance matrices

An often-used approach to identify distributed networks of fMRI signal correlations relies on a linear decomposition of the data via ICA (see below) (Beckmann et al., 2005). An alternative multivariate linear decomposition, eigenvalue decomposition, can be extended to the comparison between covariance matrices from two different conditions (e.g., drug vs. placebo). Eigenvalue decomposition identifies spatial patterns, so-called ‘spatial modes’ of correlated (or anti-correlated) signals across brain regions (Mitra and Pesaran, 1999; Friston and Büchel, 2004; Donner et al., 2013). In what follows, we first describe the standard eigenvalue decomposition (synonymous with principal component analysis) and subsequently describe its generalized form.

The eigenvalue decomposition of the AAL atlas-derived covariance matrices (C) was computed as follows:

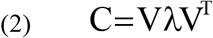

where ^T^ denoted transposition, *λ* was an *n*-by-*n* matrix with eigenvalues on its diagonal, and V was an *n*-by-*n* matrix of corresponding eigenvectors in which rows were brain regions (*n* = 90) and columns defined individual spatial modes *p*, where *p* was a vector and *p* ∈ {*p*_1_,*p*_2_,…,*p_n_*}. The overall sign of the elements in *p* was arbitrary but the sign of one element with respect to another indicated their relative covariation: equal signs indicated positive correlation and opposite signs indicated negative correlation.

For each run *i*, separately for the atomoxetine and placebo condition, we calculated participant-level time series *t* corresponding to each mode by projecting the mode onto the participant-level multivariate time series *M* via:

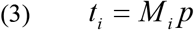

The so-computed *t* described the time-varying strength of the expression of the spatial mode (functional network) in each individual participant’s data, in one condition. We used *t* to produce voxel-level spatial maps of the corresponding modes in order to examine their correspondence with ICA-derived cofluctuating networks (see below). Next, we describe the generalization of eigenvalue decomposition to extract modes that are more strongly expressed in one condition relative to the other.

### Generalized eigenvalue decomposition of covariance matrices

We used generalized eigenvalue decomposition to decompose the covariance matrices from both experimental conditions, atomoxetine and placebo, into spatial modes that fluctuated more strongly in one condition than in the other (Friston and Büchel, 2004; Donner et al., 2013). This analysis approach has been validated for fMRI with retinotopic mapping protocols (Donner et al., 2013). Figure 1 shows a schematic overview. Using the ‘eig’ function in MATLAB 2012a, we decomposed the participant-averaged atomoxetine covariance matrix C_A_ and placebo covariance matrix C_P_ by solving the equation:

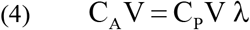

where *λ* was an *n*-by-*n* matrix with generalized eigenvalues on its diagonal, and V was an *n*-by-*n* matrix of corresponding eigenvectors in which rows were brain regions (*n* = 90 for the AAL atlas, and *n* = 140 for the Craddock atlas) and columns defined individual modes (*p*). As above, *p* was a vector and *p* ∈ {*p*_1_,*p*_2_,…,*p_n_*}. The resulting spatial modes described patterns of correlated signal fluctuations that maximized the variance (fluctuation amplitude) accounted for in one condition relative to the other (as measured by the corresponding *λ*_p_). Thus, equation 4 identified spatial modes that fluctuated more strongly in the atomoxetine condition than in the placebo condition. To identify spatial modes that fluctuated more strongly in the placebo condition, the covariance matrices C_A_ and C_P_ were swapped. We arranged V and *λ* such that their first entries corresponded to the modes that explained most variance. In other words, we sorted *λ* in descending order and then sorted V by *λ*.

**Figure 1.**
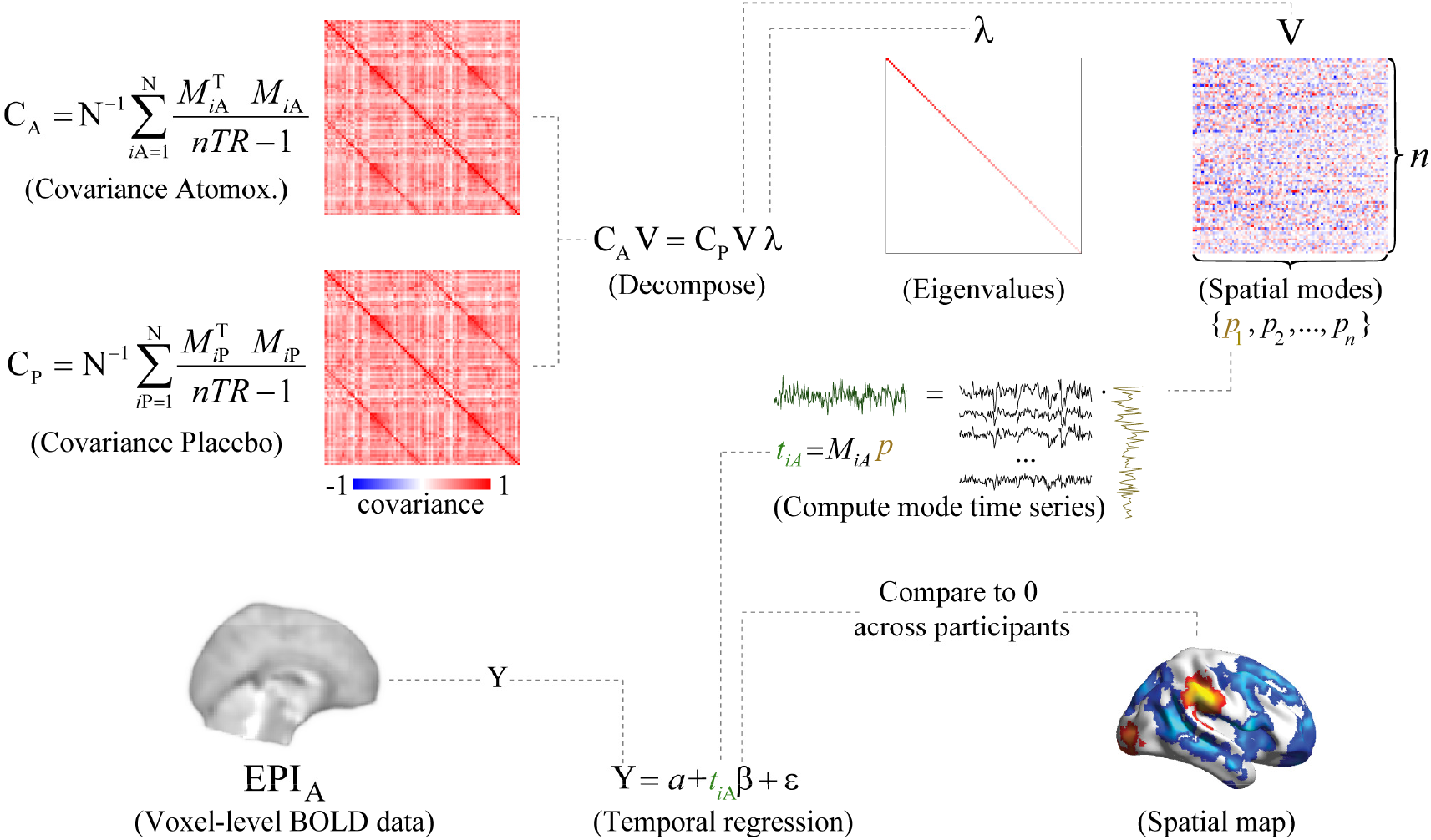
Schematic overview of the spatial mode decomposition method. The covariance matrices C_A_ and C_P_ are submitted to generalized eigenvalue decomposition to produce a matrix of eigenvalues (*λ*) and eigenvectors (V). The decomposition equation as given here delineated modes that were more strongly expressed in the atomoxetine condition than in the placebo condition. To identify modes that were more strongly expressed in the placebo condition, the covariance matrices C_A_ and C_P_ were swapped. After decomposition, the participant-level time series (*t*) corresponding to each individual spatial mode (*p*) were computed for each run *i* by projecting the mode onto the data (*M*). The number of brain regions in the parcellation scheme was denoted by *n*. A spatial map of brain regions that consistently covaried with the mode time series was computed by regressing the spatial mode time series for the atomoxetine (A) and placebo (P) conditions onto the voxel-level fMRI time series, and comparing the regression coefficients to zero across participants.

For each run *i*, we calculated participant-level time series *t_i_* corresponding to each spatial mode *p* for each individual run *i* as follows:

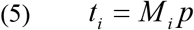

Here, *t_i_* was a vector with length 211 (the number of volumes), and *M_i_* was a matrix of Z-scored fMRI time series from the run, with size 211 by *n* (volumes by brain regions).

### Quantifying the across-subject consistency and reliability of spatial modes

The spatial modes were computed such that they explained more variance in the group-average data in the atomoxetine condition than in the placebo condition (or the converse). We aimed to quantify, in a cross-validated fashion, how consistently the fluctuation strength of these group-average spatial modes distinguished between conditions within individual subjects. The fluctuation amplitude *s_i_* corresponding to each mode’s time series in each individual run from each participant quantified the amount of variance that the mode explained in the data, and was calculated via:

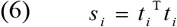

where ^T^ denoted transposition. Note that this was equivalent to:

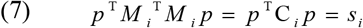

We divided *s_i_* by the sum of eigenvalues (*λ*) to convert it to units of percentage variance explained. In contrast to the eigenvalues, which captured the group-level mode’s ratio of explained variance between conditions, *s_i_* captured the amount of variance that the mode captured in the condition-specific runs at the individual participant-level. For cross-validation, we defined modes (using eq. 4) based on the group-average covariance matrices C_A_ and C_P_ that were generated from the first half of volumes in *M_i_* (using eq. 1). Each mode was projected onto independent data: the remaining half of volumes in *M_i_* as described above (eq. 5). Their corresponding fluctuation amplitudes were calculated (via eq. 6). We then used the second half of volumes to define the modes and projected them onto the first half, and averaged the two values of *s_i_*. The percentage variance explained by each mode could then be compared between conditions with non-parametric permutation testing (10,000 iterations).

We used receiver operating characteristic (ROC) analysis (Green and Swets, 1966) to quantify the reliability of the spatial modes in discriminating between experimental conditions, at the level of short segments (25% of volumes, ∼114 s) of the fMRI runs. ROC analysis performs more accurately with densely populated distributions of measurements. Thus, we defined spatial modes based on the group average covariance matrices calculated from a smaller subset of volumes (25%), as described above (using eq. 1 and eq. 4). We subdivided the remainder of volumes into 20 equal-sized bins, and computed (participant-level) *s_i_* for each of them. We cross-validated the fluctuation amplitude calculation by computing modes and projecting them onto the remaining data four times, such that eventually all data were used to define the modes. This yielded four distributions of si per condition and participant that were submitted to ROC analysis, resulting in four ROC curves per participant. We calculated the area under the ROC curve, referred to as ‘ROC index’ in the following, and averaged the resulting ROC indices across the four ROC curves of each participant. The resulting ROC indices could range between 0 and 1 and could be interpreted as the probability with which we could predict the condition from the mode’s fluctuation strength in a given data segment. The ROC indices were tested for significance by comparing them to chance level (0.5) using non-parametric permutation testing (10,000 iterations). In order to exclude the possibility that the significance of the ROC results depended on the number (25%) of volumes on which the mode was defined, we repeated the ROC analyses for modes defined on ∼14%, 20%, and ∼33% of the data, and found identical results in terms of direction and significance.

### Imaging the spatial modes

The spatial modes were computed using atlas-level covariance matrices because the whole-brain covariance matrices could not be robustly estimated at the single-voxel level (substantially more voxels in the brain than samples in the time dimension). A central aim of our study was to image the neuroanatomical distribution of the spatial modes at the single-voxel level. To this end, we used the following approach. For each participant and condition separately, we regressed the spatial mode time series *t_i_* (see eq. 5) onto the multivariate (voxel-level) time series from the corresponding run *i*. This yielded a map of regression coefficients per participant, condition, mode, and run. For each mode and for each condition, we could then compare the regression coefficients to zero using non-parametric permutation testing (10,000 iterations). The α level was set at 0.05, FWE corrected for multiple comparisons using threshold-free cluster enhancement (Smith and Nichols, 2009). The resulting statistical parametric maps indicated which voxels (if any) significantly covaried with the mode time series consistently across participants.

### Validation of spatial modes via independent component analysis

Independent component analysis (ICA) is an often-used approach to delineate so-called ‘resting-state networks’ of intrinsic fMRI signal covariations (Beckmann et al., 2005). We applied ICA in order to validate the use of eigenvalue decomposition and to examine the correspondence between spatial modes and well-characterized ‘resting-state networks’. We first estimated a set of independent components (ICs) that were representative of the combined set of resting-state runs (i.e., runs from all participants and both the atomoxetine and placebo conditions) by applying a spatial ICA to all temporally concatenated data using FSL’s MELODIC. The model order (51) was automatically estimated from the data following the methods described by Beckmann et al. (2005). Each IC represented a statistical parametric map and corresponding time series of consistent spatio-temporal dynamics. Next, we spatially correlated each IC spatial map with the 10 ‘resting-state networks’ reported by Smith et al. (2009) and selected the ICs that showed the highest correlation coefficient. The selected components showed an average correlation coefficient of 0.48 (range: 0.28 – 0.70), which indicated that the ICs as expressed in our data corresponded relatively well to previously reported ‘resting-state networks’ (Smith et al., 2009).

The 10 selected ICs were reliably expressed across the combined set of resting-state runs, and were thus representative of group-level spatiotemporal dynamics. However, the ICs did not necessarily represent spatiotemporal dynamics within individual runs. To produce a time series and a spatial map for the individual resting-state runs, we used the group-level IC spatial maps in multiple spatial regression onto the individual runs. This produced a time series for each IC as expressed within the individual runs. Then, in a second step, we used the participant-level time series as temporal regressors to produce spatial maps of regression coefficients for each component and each run. Thus, this two-stage regression approach resulted in a spatial map for each participant, condition, and IC, that indicated the degree of covariation between individual voxels and the IC time series.

To quantify the correspondence between the spatial modes and ICA-based ‘resting-state networks’, we first repeated the procedure described in the section *Imaging the Spatial Modes* above, but now on the data concatenated across the two runs runs per participant. The purpose of this concatenation procedure was to create spatial maps that were independent of the drug condition, similar to the ICs. We then correlated, across voxels, the spatial modes with the selected ICs, separately for each participant. We finally compared the distribution of Fisher *r*-to-Z transformed correlation coefficients to zero using a two-tailed *t*-test.

We also determined if standard eigenvalue decomposition identified similar spatial patterns to the more commonly used ICA. We first produced voxel-level spatial maps of the modes that were derived from eigenvalue decomposition of AAL atlas-level covariance in the individual conditions, using multiple temporal regression. We then selected modes based on maximal spatial correlation with the 10 intrinsic connectivity networks reported by Smith et al. (2009), similar to the selection of ICA components described above. Finally, we examined the strength of correlation between the selected voxel-level spatial mode maps and the intrinsic connectivity networks reported by Smith et al. (2009).

### Similarity between spatial modes from different parcellation schemes

We used spatial correlation to determine if the generalized eigenvalue decomposition-derived mode spatial maps depended on the parcellation scheme. For each individual participant and condition, we correlated the (unthresholded) spatial maps of regression coefficients of the modes that were generated with the AAL atlas, and those that were generated with the Craddock atlas. We then compared the distribution of Fisher-transformed correlation coefficients to zero using a two-tailed *t*-test. Similarly, we characterized the correspondence in mode spatial maps between the individual conditions by correlating the unthresholded spatial maps at the individual participant level, and comparing the resulting distribution of Fisher-transformed correlation coefficients to zero using a two-tailed *t*-test.

### Similarity between spatial modes and catecholamine receptor expression maps

We used a dataset provided by the Allen Brain Institute (Hawrylycz et al., 2012; Hawrylycz et al., 2015) (http://www.brain-map.org/) to quantify the similarity between the spatial modes (computed based on signal fluctuations as described above) and the spatial maps of the expression of specific catecholamine receptors. The Hawrylycz et al. (2015) dataset comprised post-mortem samples of 6 individuals that underwent microarray transcriptional profiling. Spatial maps of each sample’s gene transcription profile were available in MNI space, following improved non-linear registration as implemented by Gorgolewski et al. (2014). Receptors mediate the effect of neuromodulators on post-synaptic neurons and, consequently, neural network dynamics. In the current article we thus focused on the expression of clusters of genes that encode receptors with varying subunit compositions but functionally analogous post-synaptic effects (e.g. due to being coupled to inhibitory or excitatory G-proteins). Specifically, we grouped the 14 available catecholamine receptor-related genes into 5 classes according to functional receptor type: norepinephrine receptor α_1_ (ADRA1A, ADRA1B, ADRA1D); norepinephrine receptor α_2_ (ADRA2A, ADRA2B, ADRA2C); norepinephrine receptor β (ADRB1, ADRB2, ADRB3); and dopamine ‘D_1_- like’ (DRD1, DRD5) and ‘D_2_- like’ (DRD2, DRD3, DRD4) receptors (Cools and van Rossum, 1976; Surmeier et al., 2007; Arnsten, 2011).

We used two groups of “reference” receptors in order to examine the specificity of the spatial similarity measures for catecholamine receptors. First, because the cholinergic system has a gross functional organization similar to the norepinephrinergic system (e.g., cortex-wide cholinergic projections), we used an additional 16 genes related to acetylcholine receptors as a reference. Those were grouped into two classes, again according to functional receptor type: nicotinic acetylcholine receptor (ACh_N_ (CHRNA2, CHRNA3, CHRNA4, CHRNA5, CHRNA6, CHRNA7, CHRNA9, CHRNA10, CHRNB2, CHRNB3, CHRNB4) and muscarinic acetylcholine receptor (ACh_M_) (CHRM1, CHRM2, CHRM3, CHRM4, CHRM5). Second, because atomoxetine also blocks NMDA receptors (Ludolph et al., 2010), we selected 7 genes related to the expression of NMDA receptors (GRIN1, GRIN2A, GRIN2B, GRIN2C, GRIN2D, GRIN3A, GRIN3B). These genes were grouped into one class because NMDA receptor blockade by atomoxetine is similar across receptors with varying subunit compositions (Ludolph et al., 2010).

Spatial similarity (i.e., correlation) between gene expression and spatial modes (imaged at the single-voxel level, see above) was computed on an individual participant basis by linear regression across sequenced parcels. Because the post-mortem samples differed in the coverage of sequenced parcels, we repeated this procedure for each individual post-mortem sample. A *t*-test was then conducted across samples to obtain a test statistic that quantified the robustness of the spatial correlation across the 6 post-mortem samples (Gorgolewski et al., 2014). For each of our participants, we then collapsed across genes within each receptor class (α_1_, α_2_, β, D_1_-like, D_2_-like, ACh_N_, ACh_M_, and NMDA). To assess the robustness of correlations across our participants (in addition to within participants across samples), we compared the distribution of *t*-statistics of the catecholamine receptors to zero, and to the *t*-statistics of the acetylcholine receptors and NMDA receptors, by means of non-parametric permutation testing (10,000 iterations). Significant differences of *t*-values were indicative of a relationship between the expression of specific catecholamine receptors and the spatial distribution of the modes that was reliable across both post-mortem samples and across our participants.

Acetylcholine receptors were unrelated to the spatial mode maps, and Bayes factors indicated “substantial” evidence (Wetzels and Wagenmakers, 2012) for the null hypothesis of no correlation (ACh_N_ and spatial mode atomoxetine > placebo: *p* = 0.28, BF = 0.157; for ACh_N_ and spatial mode placebo > atomoxetine: *p* = 0.15, BF = 0.156; for ACh_M_ and spatial mode atomoxetine > placebo: *p* = 0.26, BF = 0.157; for ACh_M_ and spatial mode placebo > atomoxetine: *p* = 0.73, BF = 0.157).

Moreover, there were no significant differences between the muscarinic and nicotinic acetylcholine receptors (spatial mode atomoxetine > placebo versus spatial mode placebo > atomoxetine, ACh_N_,: *p* = 0.07; ACh_M_, *p* = 0.31). We thus collapsed across acetylcholine receptors and used this summary statistic as reference for testing the mode versus receptor map associations for the catecholamine receptors. Similarly, we found no significant associations between NMDA receptors and spatial modes, and Bayes factors indicated “substantial” evidence for the absence of a correlation (atomoxetine > placebo: *p* = 0.81, BF = 0.157; placebo > atomoxetine: *p* = 0.11, BF = 0.156), and thus used NMDA receptors as an additional reference.

### Separating spatial modes from noise

We calculated the theoretical distribution ρ of eigenvalues *λ* under the null hypothesis of no difference between conditions, and was given by the following:

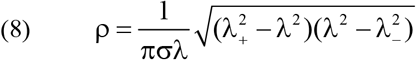

where:

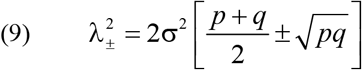

and σ was the standard deviation of *λ*, and p and q were the dimensions of the covariance matrix. We then fitted ρ to *λ* by minimizing the sum of squared residuals of ρ multiplied by a scalar value (Mitra and Pesaran, 1999).

If between-condition differences in signal correlation strength were “noise” (i.e., independently normally distributed with zero mean), the eigenvalues should not have differed from the theoretical distribution (Mitra and Pesaran, 1999). If, by contrast, the between-condition differences in correlation strength were “signal”, the eigenvalues of modes with a low rank number should have exceeded the theoretical distribution more so than modes with a high rank number, reflecting a skewed eigenvalue distribution. We thus calculated the difference between the eigenvalues and the theoretical distribution and categorized modes into “signal” and “noise”. Modes for which *λ* > ρ were categorized as signal; the remaining modes were categorized as noise. This procedure provided an upper bound for the number of modes that we could consider as possibly reflecting atomoxetine-related changes in intrinsic signal correlation strength.

### Control analyses for importance of first spatial mode

We performed two control analyses to examine to what extent the first spatial mode captured atomoxetine effects (on the strength of correlations in relation to catecholamine receptors) over and above the subsequent spatial modes classified as signal (see previous section). First, we determined if all signal modes with a lower rank number tended to explain more variance in independent data than modes with a high rank number, using the cross-validated ROC analysis described above. If so, the ROC index should decline with mode rank. Note that this prediction was not trivial given that in the cross-validation procedure the modes were projected onto independent data. We tested this prediction by correlating ROC index with mode rank within participants and comparing the distribution of correlation coefficients to zero across participants using permutation testing (10,000 iterations, one-tailed test).

Second, we determined if the spatial correspondence between modes and catecholamine receptors was stronger for the first mode than for the subsequent signal modes (i.e., rank numbers > 1). We used permutation tests to compare the corresponding spatial correlations between mode one and the remaining modes: once by collapsing correlations across signal modes and once for all subsequent modes individually.

### Control analysis for mode specificity

Spatial mode decomposition (eqn. 4) can only be used to compare two individual conditions (or groups): here, the placebo and atomoxetine conditions. However, the fMRI measurements of the atomoxetine and placebo conditions were conducted on separate days. Thus, it is possible that spatial modes reflected session-related effects rather than drug treatment-related effects. To control for this possibility, we projected the spatial modes onto the multivariate fMRI data (using eqn. 5) of the pre-pill measurements that were conducted on the same days as the post-pill ingestion measurements, and calculated the strength of the fluctuation of the resulting time series (using eqn 6.). We then used the percentage of variance explained in the pre-pill measurements as a baseline in the interaction contrast (atomoxetine - pre atomoxetine) - (placebo - pre placebo).

Second, we computed spatial modes based on covariance in the pre-pill ingestion conditions, and compared (using spatial correlation) the resulting spatial maps to those that were computed using the post-pill measurements. We then compared the distribution of correlation coefficients across participants to zero using permutation testing.

### Code availability

MATLAB code to compute spatial modes and run statistical analyses of mode variance can be found here ([link inserted upon acceptance]).

## RESULTS

The aim of the present study was to assess the spatial distribution of catecholaminergic modulation of large-scale brain dynamics and relate it to the spatial distribution of catecholamine receptors. To this end, we imaged atomoxetine-induced alterations (increases and decreases) in the strength of correlated fMRI signal fluctuations across the whole human brain, and related the resulting spatial maps (referred to as ‘spatial modes’, see Materials and Methods) to maps of catechacholamine receptor gene expression derived from post-mortem brains (Hawrylycz et al., 2012; Hawrylycz et al., 2015). We used a linear decomposition approach, which we previously validated by means of fMRI retinotopic mapping protocols (Donner et al., 2013), and which was tailored to finding the two spatial modes that fluctuated more strongly (referred to as ‘atomoxetine > placebo’) or less strongly (‘placebo > atomoxetine’) during the atomoxetine condition than during the placebo condition (Figure 1 and Materials and Methods). This analysis enabled imaging the brain-wide distribution of the strongest catecholamine-induced increases and decreases in correlated signal fluctuations, thus assessing their fine-grained neuroanatomical distribution. Furthermore, the analysis enabled us to quantify the similarity between the spatial modes that captured catecholaminergic modulation of brain dynamics on the one hand, and the spatial distribution of the expression of specific catecholamine receptor genes on the other hand. The latter was taken from a dataset provided by the Allen Brain Institute (Hawrylycz et al., 2012; Hawrylycz et al., 2015).

The Results section is organized as follows. We first describe the spatial modes that show the strongest drug-induced changes (increases and decreases) in correlated signal fluctuations. We then evaluate the relationship between both of these spatial modes and catecholamine receptor gene expression maps. Finally, we present a number of control analyses that support the specificity and validity of the spatial modes of drug-related changes in brain dynamics.

### Spatial modes fluctuating more strongly during atomoxetine than placebo

Our previously published analyses of the same data (van den Brink et al., 2016) identified only *reductions* in strength of inter-regional fMRI signal correlations. Our current approach uncovered a distributed pattern (i.e., spatial mode) of correlated signal fluctuations that *increased* under atomoxetine (Figure 2). The eigenvalues all 90 spatial modes (as many as brain regions in the AAL atlas) for the atomoxetine > placebo comparison are shown in Figure 2a. Here, we focused on analyzing the first of these spatial modes (Figure 2b) because it had the largest eigenvalue, thus exhibiting the strongest increase in fluctuation amplitude during the atomoxetine condition, and because mode orthogonality can obscure the interpretation of modes with higher ranks (c.f. Donner et al., 2013).

**Figure 2.**
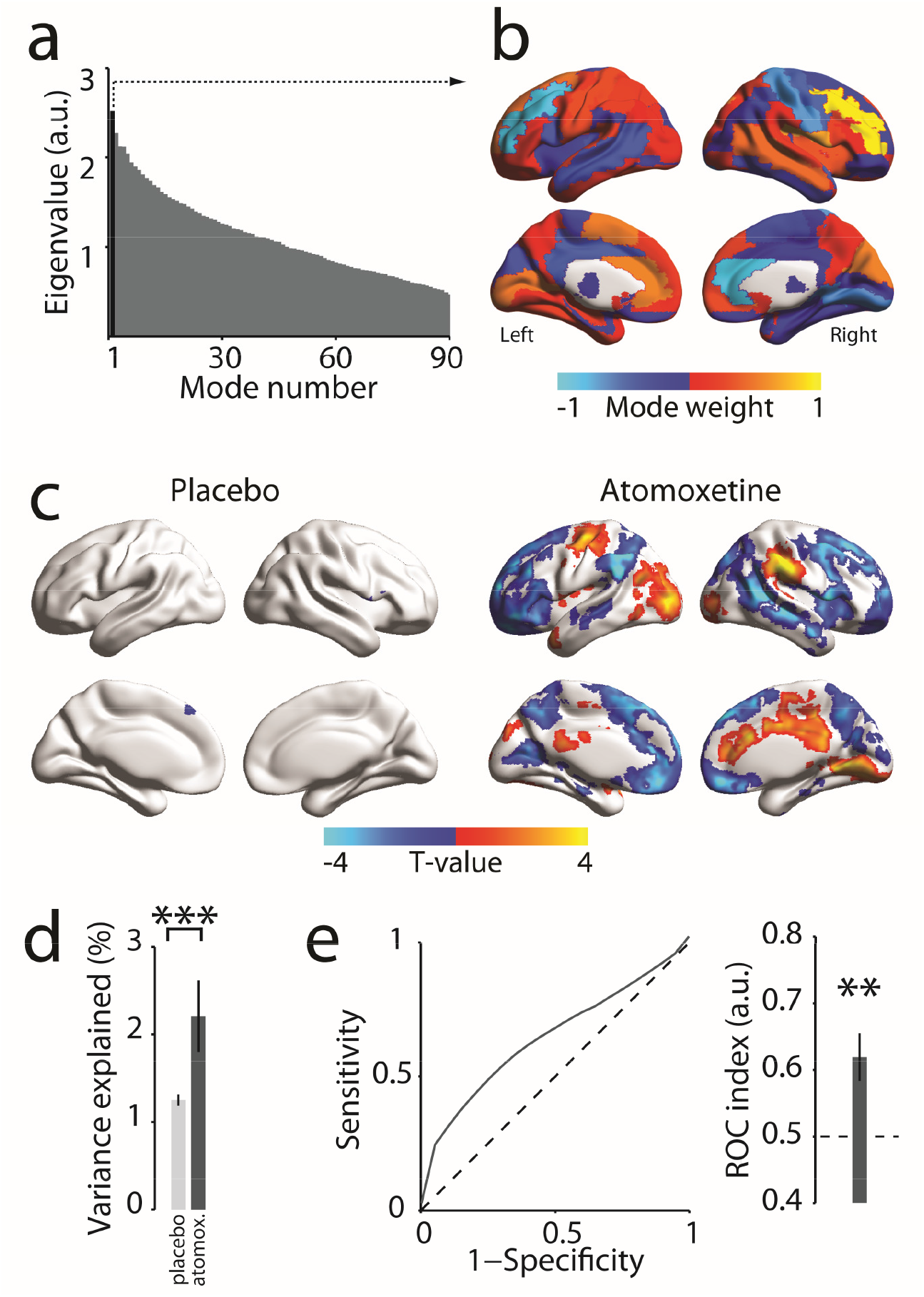
Spatial mode that fluctuated more strongly during atomoxetine than placebo. ***a*,** Eigenvalue spectrum of all spatial modes from generalized eigenvalue decomposition of covariance matrices for atomoxetine > placebo (AAL atlas, see Materials and Methods). Black, eigenvalue of the first spatial mode. ***b*,** Distribution of spatial mode weights visualized on cortical surface reconstruction. ***c*,** Voxel-level map of significant expression of first spatial mode per condition (*p* < 0.05, FWE corrected). ***d*,** Comparison of percentage of variance explained by spatial mode from panel *b* in the atomoxetine and placebo conditions. Independent data were used for computing the spatial mode and assessing the variance explained. ***e*,** ROC-analysis of discriminability of conditions based on fluctuation amplitude of the first spatial mode within short data segment (∼114 s, again independent of data used for computing the mode; see Materials and Methods). ROC indices > 0.5 indicate that the spatial mode fluctuation predicts the condition. Error bars, SEM across participants (N=24). **,*p* < 0.01; ***,*p* < 0.001.

The spatial mode was comprised of a set of weights (one value per brain region in the parcellation scheme) that indicated relative cofluctuation between brain areas (Figure 2b). Please note that the overall sign of mode weights was arbitrary, but the sign of one element with respect to another indicated their relative phase, with equal signs indicating positive correlation and unequal signs indicating negative correlation. The mode displayed maxima (both positive and negative) in bilateral middle frontal gyri, bilateral anterior cingulate cortices, right lingual gyrus and postcentral gyrus, left calcarine fissure and surrounding cortex, and in the left supplementary motor area. Across the brain, the weights were anti-correlated between hemispheres (*r* = -0.56, *p* < 0.001) such that if the mode weight of one brain region was positive then the weight of the homotopic region in the other hemisphere tended to be negative. This suggests that this spatial mode possibly reflected an increase in the mutual inhibition between hemispheres.

The spatial mode shown in Figure 2b was a coarse (atlas-level) representation of the spatial distribution of the corresponding brain dynamics, which was necessary for technical reasons (Materials and Methods). Regressing the time series of the fluctuation of this spatial mode onto each participant’s multivariate data enabled us to image this spatial distribution at a finer (voxel-level) granularity, as well as test for the consistency of the expression of the corresponding spatial mode across participants within individual conditions (Figure 2c). This analysis yielded a single significant cluster (superior frontal gyrus) for the placebo condition, and a number of significant clusters (36% of all brain voxels) for the atomoxetine condition. Cluster maxima were located in bilateral anterior cingulate cortex, right medial frontal gyrus, right lingual gyrus, left precentral gyrus, left lateral occipital cortex, and bilateral supramarginal gyri.

To determine if the spatial mode corresponded to any of the so-called ‘resting-state networks’, as defined with commonly used ICA approaches (Beckmann, 2009; Smith et al., 2009), we correlated the spatial mode with each of the 10 selected ICA components (see Materials and Methods) for each individual participant. This yielded a weak, albeit statistically significant, correlation of the spatial mode with the right-lateralized frontoparietal ICA component (mean *r* = -0.05, SD 0.03; *t*(23) = -7.89, *p* < 0.001). Taken together, our analysis revealed a pattern of intrinsic fMRI signal correlations that were enhanced under atomoxetine, which exhibited a highly structured spatial organization (Figure 2c), but only loosely resembled any of the established resting-state networks defined using standard ICA-based analyses of correlated signal fluctuations irrespective of pharmacological intervention. In the section *Spatial Modes Reflect Gene Expression of Catecholamine Receptors*, we link this spatial organization to the distribution of specific catecholamine receptors across the brain.

We finally verified, using cross-validated procedures (Materials and Methods), the robustness and reliability of the fluctuations captured by the spatial mode: The fluctuation strength of the spatial mode was consistently larger in the atomoxetine than placebo condition (*p* < 0.001; Figure 2d), and it reliably discriminated between the two pharmacological conditions, even on the basis of short individual data segments (group average ROC index = 0.62, *p* = 0.002; Figure 2e).

### Spatial modes fluctuating less strongly during atomoxetine than placebo

While our previous work identified catecholamine-related reductions in the overall strength of correlated signal fluctuations (van den Brink et al., 2016), the analysis approach we used previously was not suited to image the fine-grained neuroanatomical structure of these decreases. By contrast, our current decomposition approach suited this purpose, and it uncovered a widespread set of brain regions between which correlations were suppressed by atomoxetine (Figure 3). The first spatial mode resulting from this decomposition (again selected based on its largest eigenvalue, Figure 3a) had local maxima and minima in homotopic regions of both hemispheres (Figure 3b), with an even stronger overall negative correlation between hemispheres (*r* = -0.79, *p* < 0.001) than evident for the spatial mode for atomoxetine-induced increases (compare to Figure 2b). This effect might indicate an catecholamine-induced reduction in inter-hemispheric competition.

**Figure 3.**
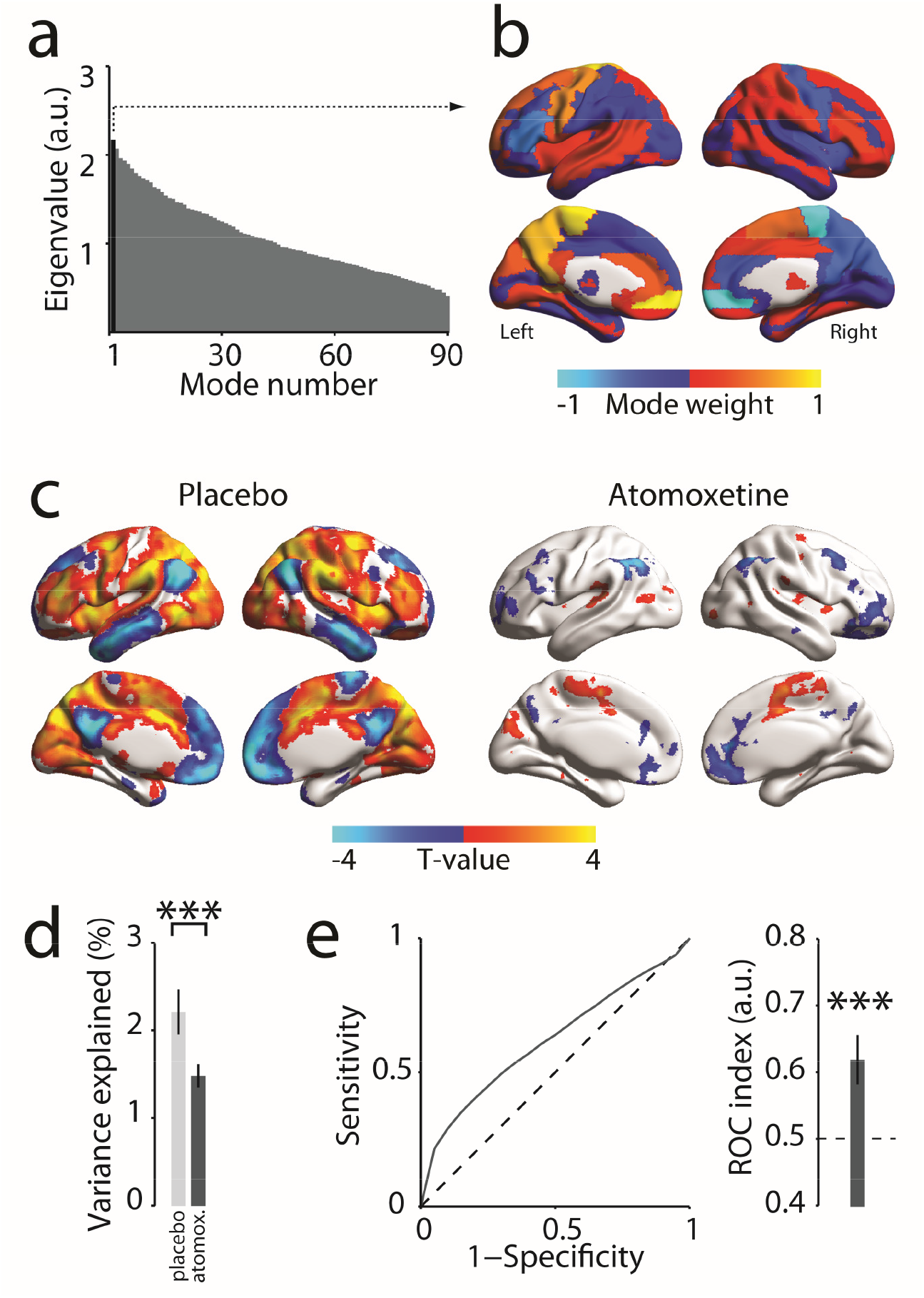
Spatial mode that fluctuated less strongly during atomoxetine than placebo. ***a*,** Eigenvalue spectrum of all spatial modes from generalized eigenvalue decomposition of covariance matrices for placebo > atomoxetine. Black, eigenvalue of the first spatial mode. ***b*,** Distribution of spatial mode weights visualized on cortical surface reconstruction. ***c*,** Voxel-level map of significant expression of first spatial mode per condition (*p* < 0.05, FWE corrected). ***d*,** Comparison of percentage of variance explained by spatial mode in (panel *b*) in the atomoxetine and placebo conditions. ***e*,** ROC-analysis of discriminability of conditions based on fluctuation amplitude of the first spatial mode within short data segment Error bars, SEM across participants (N=24). ***, *p* < 0.001.

Importantly, this spatial mode for placebo > atomoxetine was uncorrelated (*r* = -0.013, *p* = 0.88, Bayes Factor = 0.157) with the one for atomoxetine > placebo (Figure 2b). Thus, the spatial modes resulting from both decompositions reflected distinct sets of brain regions, in which the direction of catecholaminergic effects on signal correlation strength was opposite.

Again, we imaged the fine-grained (voxel-level) distribution of this fluctuation pattern within individual conditions. This revealed a large proportion of significant voxels (51% of all brain voxels) in the placebo condition (Figure 3e). The spatial mode exhibited local maxima or minima in regions of the so-called ‘default mode’ and ‘attention networks’ (Fox et al., 2006; Smith et al., 2009): bilateral temporal poles, medial frontal, lateral occipital, and posterior cingulate cortices, and in bilateral paracingulate, precentral, superior frontal, supramarginal, and paracingulate gyri. Indeed, the spatial mode weakly, but significantly and most strongly, resembled the left-lateralized ‘frontoparietal’ ICA component (mean *r* = -0.15, SD 0.05; *t*(23) = -16.33, *p* < 0.001). Given that the first spatial mode in the decomposition atomoxetine > placebo correlated most strongly with the right lateralized frontoparietal network, this suggested that atomoxetine resulted in a shift from left-to right-lateralized frontoparietal dominance. A significant interaction in the strength of correlation between mode polarity (atomoxetine-induced increase versus decrease) and ICA component (frontoparietal left versus right) suggested that this was indeed the case (repeated-measures ANOVA; *F*(1,23) = 163.14, *p* < 0.001). Other significant correlations were evident for the ‘default mode’ (mean *r* = -0.15, SD 0.05; *t*(23) = -14.19, *p* < 0001) and ‘sensorimotor’ (mean *r* = 0.13, SD 0.04; *t*(23) = 17.41, *p* < 0.001) ICA components.

The fluctuation of this spatial mode was a consistent and reliable indicator of the drug condition across participants, even for short segments of data (comparison of mode variance between atomoxetine and placebo: *p* < 0.001; group average ROC index = 0.62, *p* = 0.002; Figure 3d,e).

### Spatial modes reflect gene expression of catecholamine receptors

Our analyses thus far established that atomoxetine both increased and decreased intrinsic fMRI signal correlations in two distinct sets of widely distributed brain regions. How can the systemic increase in catecholamine levels by atomoxetine lead to regionally specific, and even opposite-polarity modulations of brain dynamics? An attractive possibility is that such heterogeneous functional effects are mediated by the heterogeneous distribution of catecholamine receptors across the brain (Ramos and Arnsten, 2007). To test this idea, we quantified the spatial similarity between spatial modes and maps of the expression of genes encoding a variety of catecholamine receptors.

Gene maps were taken from human post-mortem samples from the Allen Brain Institute (Hawrylycz et al., 2012; Hawrylycz et al., 2015) and examples are shown in Figure 4a. We found a specific, and distinct, association pattern for both spatial modes identified here (Figure 4b). First, the spatial mode that fluctuated more strongly in the atomoxetine than the placebo condition was associated with the genetic expression map of D_2_-like dopamine receptors. Second, by contrast, the spatial mode that fluctuated more strongly in the placebo than atomoxetine condition was associated with genetic expression of the α_1_ norepinephrine receptor. Third, both spatial modes were associated with the β norepinephrine receptor gene map, but with opposite sign.

**Figure 4.**
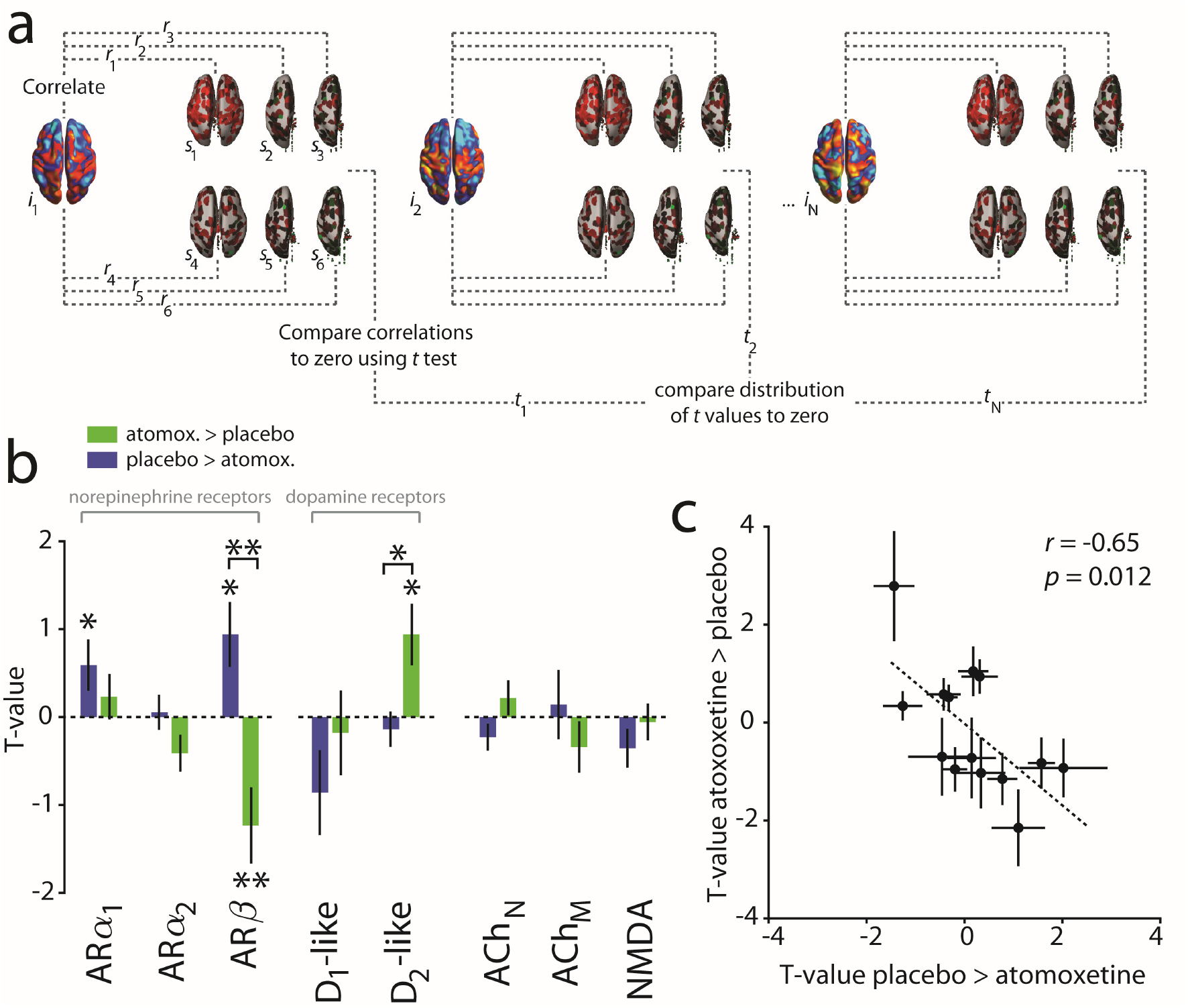
Associations between spatial modes and catecholamine receptor gene expression. ***a*,** Correlation between individual spatial modes (*i*) and 6 post-mortem samples (*s*). This procedure was repeated for each gene. See Materials and Methods for details. ***b*,** Correlations between spatial modes and receptor gene expression maps. Within-mode significance is assessed by comparison to zero. AR, adrenoceptor; ACh, acetylcholine receptor. *, *p* < 0.05; **, *p* < 0.01. ***c*,** Relationship of spatial mode vs. catecholamine gene associations between both spatial modes. Error bars, SEM (N=24 participants).

To assess the specificity of these spatial correlations (Materials and Methods) we compared them with two “reference” correlations: (i) correlations with maps of genes coding for acetylcholine receptors, and (ii) correlations with maps of genes coding for NMDA receptors. We chose acetylcholine because it is another neuromodulaory system with a functional organization similar to that of the NE system, but had no relation to our drug manipulation. We chose NMDA receptors because atomoxetine binds to, and inhibits, them at clinically relevant doses (Ludolph et al., 2010). The distributions of acetylcholine receptors and NMDA were uncorrelated with the spatial mode maps (all Bayes Factors: 0 < BF < 0.158; see Materials and Methods), and there were no significant differences between the muscarinic and nicotinic acetylcholine receptors (Figure 4b, rightmost panel and Materials and Methods for details).

All the significant associations between spatial modes and catecholamine receptors shown in Figure 4b were also significant when compared to ACh receptor maps combined or to NMDA receptor maps (Comparison with ACh receptors: spatial mode atomoxetine > placebo: ARβ, *p* = 0.011; D_2_-like receptors, *p* = 0.009; spatial mode placebo > atomoxetine: ARα_1_, *p* = 0.011; ARβ, *p* = 0.010; Comparison with NMDA receptors: spatial mode atomoxetine > placebo: ARβ, *p* = 0.026; D_2_- like receptors, *p* = 0.003; spatial mode placebo > atomoxetine: ARα_1_, *p* = 0.002; ARβ, *p* = 0.003). Furthermore, similar results were obtained when the analysis was confined to the cortex rather than the whole brain, except that the association between the β norepinephrine receptor map and the spatial modes were no longer significant (spatial mode for placebo > atomoxetine: *p* = 0.07; spatial mode for atomoxetine > placebo: *p* = 0.422), but only the difference between these two associations was significant (difference between modes: *p* = 0.049). These latter findings are consistent with the relatively high expression of β receptors in subcortical areas compared to cortical areas (Rainbow et al., 1984; Reznikoff et al., 1986; Joyce et al., 1992; van Waarde et al., 1997).

The spatial mode / gene map associations were negatively correlated between the two spatial modes assessed here (Figure 4c). In other words, the more similar (dissimilar) the spatial distribution of a particular catecholamine receptor gene was to the spatial mode that showed the atomoxetine-induced *increase* in fluctuations, the more dissimilar (similar) this distribution was to the spatial mode that showed the atomoxetine-induced *reduction* in fluctuations. This was despite the fact that the spatial modes per se were unrelated to one another (see above, section *Spatial Modes Fluctuating Less Strongly during Atomoxetine than Placebo*).

In sum, the spatial association analyses reported here link the brain-wide distribution of catecholaminergic effects on large-scale neural dynamics to the distribution of different catecholamine receptor types, with important implications for understanding the principles of cathecholaminergic modulation (see Discussion). In the remainder of Results section, we present a number of control analyses, which corroborated the specificity and validity of the interpretation of our main findings.

### Control 1: Mode 1 uniquely captures atomoxetine-related effects on correlations in relation to specific catecholamine receptors

It is possible that spatial modes other than the first mode we focused on here, captured meaningful relationships between the spatial distributions of catecholamine receptors and the distribution of atomoxetine-related changes in fMRI signal correlations. To assess the relevance of spatial modes with higher ranks, we computed a theoretical distribution of eigenvalues under the null hypotheses of no between-condition differences in correlations, and compared it to the observed eigenvalue distribution (see Materials and Methods). While for both decomposition directions the first mode was clearly discernible in its deviance from the theoretical distribution, a number of subsequent modes also reflected signal (21 modes in Figure 5a, 26 modes in Figure 5c). Yet, two observations indicated that the first spatial modes (for both decomposition directions) captured the predominant effects of atomoxetine. First, they tended to explained a larger proportion of variance in one condition relative to another than the remaining ones: for both decomposition directions, the ROC index was strongly negatively correlated with mode rank number (Figure 5b,d). Second, the first spatial mode exhibited significantly stronger correlations with the distributions of catecholamine receptors than the subsequent signal modes (Figure 5e,f), and no individual signal mode correlated more strongly with catecholamine receptors than the first mode (smallest corrected p-values and Bayes Factors: atomoxetine > placebo: *p* = 0.13, BF = 0.93; placebo > atomoxetine: *p* = 0.28, BF = 0.75). Thus, the first mode optimally reflected atomoxetine-induced changes in signal correlation strength in relation to the distribution of specific catecholamine receptors.

**Figure 5.**
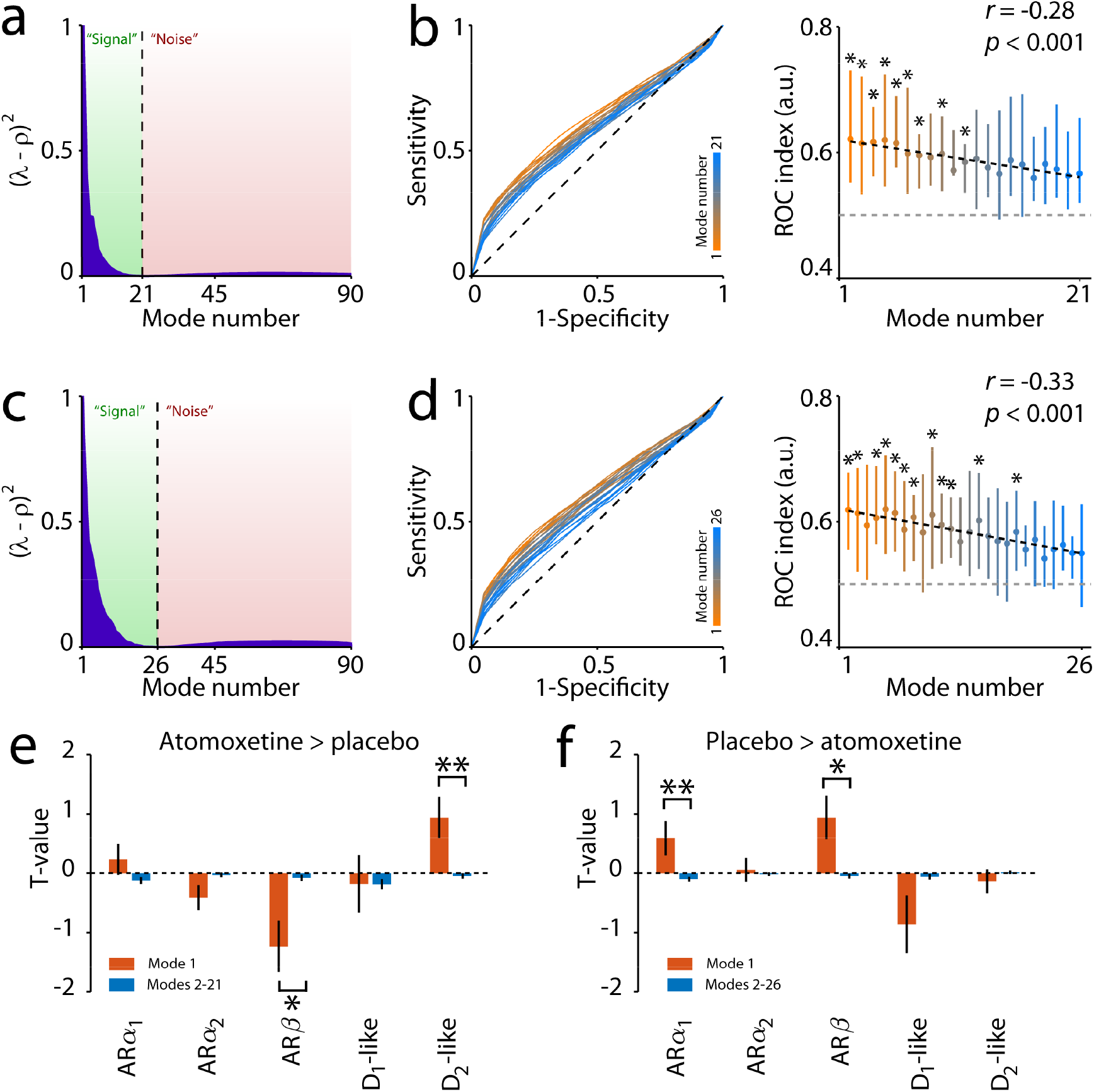
Results of control analyses for mode selection. ***a*,** Atomoxetine > placebo: difference between theoretical “noise” distribution ρ and eigenvalues. ***b***, Atomoxetine> placebo: ROC curves and indices for all modes that were categorized as “signal”. *:FDR *p* < 0.05. Error bars show a 66% CI. ***c***,***d*** same as *a* and *b* but for the decomposition direction placebo > atomoxetine. ***e***,***f*** Correlations between modes and receptor gene expression. *: *p* < 0.05; **: *p* < 0.01.

### Control 2: Spatial modes reflect drug-induced, not session-related, differences in signal fluctuations

The linear decomposition analysis performed here, by design, returned a spatial mode of which the fluctuation strength differed between the two conditions that were used to calculate the spatial mode (here: atomoxetine and placebo). Our reliability analyses established that the two spatial modes shown in Figures 2b and 3b accurately discriminated between pharmacological conditions, even in short stretches of data independent from the ones used to identify the modes (Figures 2d,e and 3d,e). This establishes that both spatial modes captured meaningful alterations of brain dynamics, rather than measurement noise. Nevertheless, they may have reflected changes in brain dynamics that differed systematically between the placebo and atomoxetine sessions, without reflecting specific drug treatment effects. Specifically, because both sessions took place one week apart, it was possible that the spatial modes might have reflected the session rather than the treatment.

We addressed this concern by analyzing the pre-pill ingestion fMRI measurements that took place on the same days. We projected the spatial modes onto the multivariate fMRI data of the pre-pill measurements and calculated the strength of the fluctuation of the resulting time series (variance explained, see eqn. 6 in the Materials and Methods). If the spatial modes reflected changes in brain dynamics that were specifically due to the catecholaminergic intervention rather than to session differences, then (i) their fluctuation amplitudes should differ more for the post-pill measurements than for the pre-pill measurements, and (ii) spatial modes computed in an analogous fashion for the pre-pill ingestion conditions should exhibit a different spatial structure from the spatial modes we investigated so far. That is what we found (Figure 6). First, the interaction contrast (atomoxetine - pre atomoxetine) - (placebo - pre placebo) was significant, in the expected direction for both spatial modes analyzed here (Figure 6a). Second, the spatial modes that were computed for the pre-pill measurements (Figure 6b) did not resemble those from the post-pill measurements (compare with Figures 2b and 3b), with no significant spatial correlations (all absolute r values < 0.06, all *p* values > 0.60). Taken together, these control analyses rule out session-related effects as a confound and further establish that the spatial modes assessed in the previous sections reflected drug-induced changes in brain dynamics.

**Figure 6.**
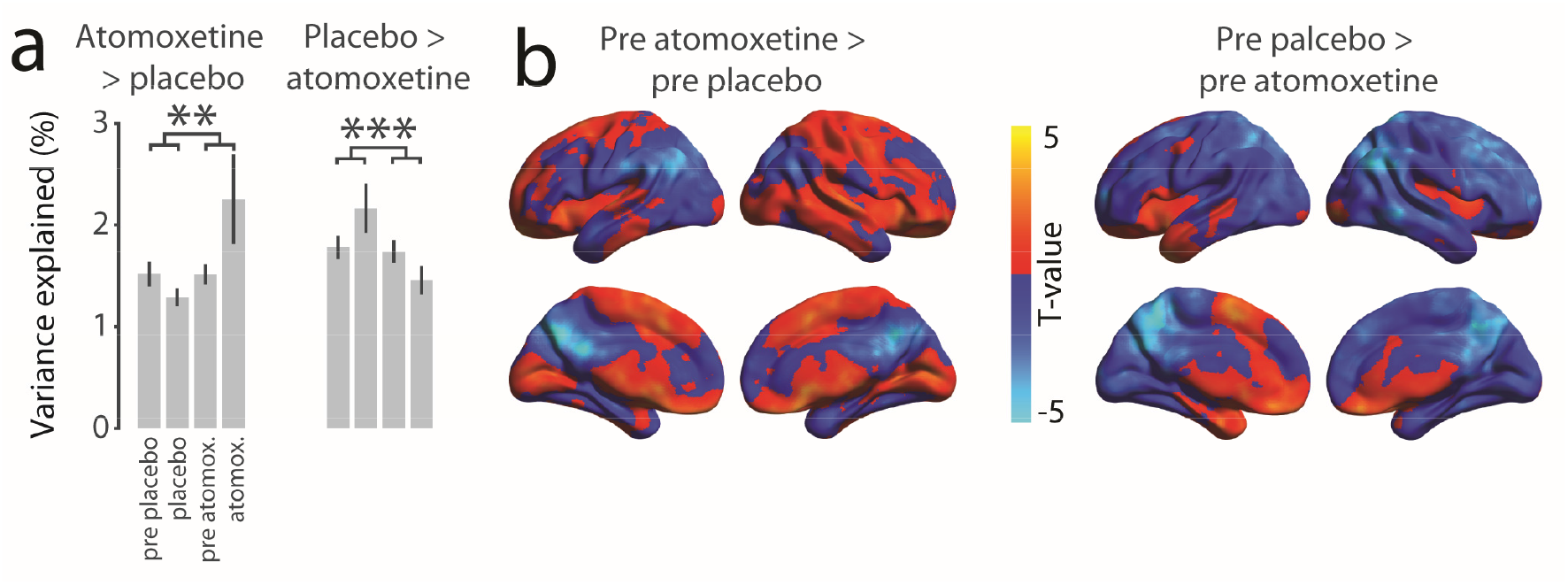
Results of control analyses examining mode specificity. ***a*,** Percentage of variance explained by the AAL atlas-derived first modes in relation to the percentage of variance explained when the modes were projected onto the pre pill ingestion measurements. Error bars, SEM across participants (N=24). Asterisks denote interaction effects of the contrast (atomoxetine vs. pre atomoxetine) vs (placebo vs pre placebo): **: *p* < 0.01; ***: *p* < 0.001. ***b*,** Threshold-free spatial maps of AAL atlas-derived modes that were generated using only the pre pill ingestion conditions. Maps are shown only for the pre placebo condition for brevity. Colored regions show covariation with the mode time series.

### Control 3: Craddock parcellation yields similar results as AAL parcellation

In order to rule out that our results depended on the specific anatomical parcellation scheme used for computing the spatial modes (AAL), we repeated the analyses using an alternate atlas that resulted from a functional parcellation and had a higher density (Craddock et al., 2012). Both resulting spatial modes explained more variance in one condition than in the other, in the expected direction (atomoxetine > placebo: *p* < 0.001; placebo > atomoxetine: *p* < 0.001). Again, these effects were reliable at the level of independent and short (∼114 s) data segments (atomoxetine > placebo: *p* < 0.001; placebo > atomoxetine: *p* < 0.001). Thus, the Craddock parcellation also yielded spatial modes that reliably differed between the two pharmacological conditions in terms of fluctuation strength.

The resulting spatial modes were also similar to the ones from our main analyses in terms of their spatial structure. To establish this, we again imaged the expression of the spatial mode time series across all brain voxels and compared the resulting map to the corresponding map from the AAL parcellation in our main analyses (Figure 7, Figure 8). Despite using parcellation schemes that differed both in the number of brain regions and in the way the brain regions were defined (anatomical parcellation and functional clustering, respectively), the mode spatial maps generated with the two atlases corresponded robustly across participants for the spatial mode atomoxetine > placebo (placebo: *t*(23) = 3.96, *p* < 0.001; atomoxetine: *t*(23) = 3.98, *p* < 0.001, Figure 7). Moreover, the spatial modes, imaged at single-voxel level, correlated between drug conditions (AAL atlas: *t*(23) = 6.93, *p* < 0.001; Craddock atlas: *t*(23) = 14.89, *p* < 0.001; Figure 7). This was also the case for the spatial mode placebo > atomoxetine: spatial modes correlated across atlases (placebo: *t*(23) = 10.43, *p* < 0.001; atomoxetine: *t*(23) = 9.54, *p* < 0.001; Figure 8) and drug conditions (AAL: *t*(23) = 15.57, *p* < 0.001; Craddock: *t*(23) = 14.89, *p* < 0.001; Figure 8). In sum, the Craddock atlas-derived modes yielded similar results in terms of direction and significance of effects as well as spatial structure of the resulting spatial modes.

**Figure 7.**
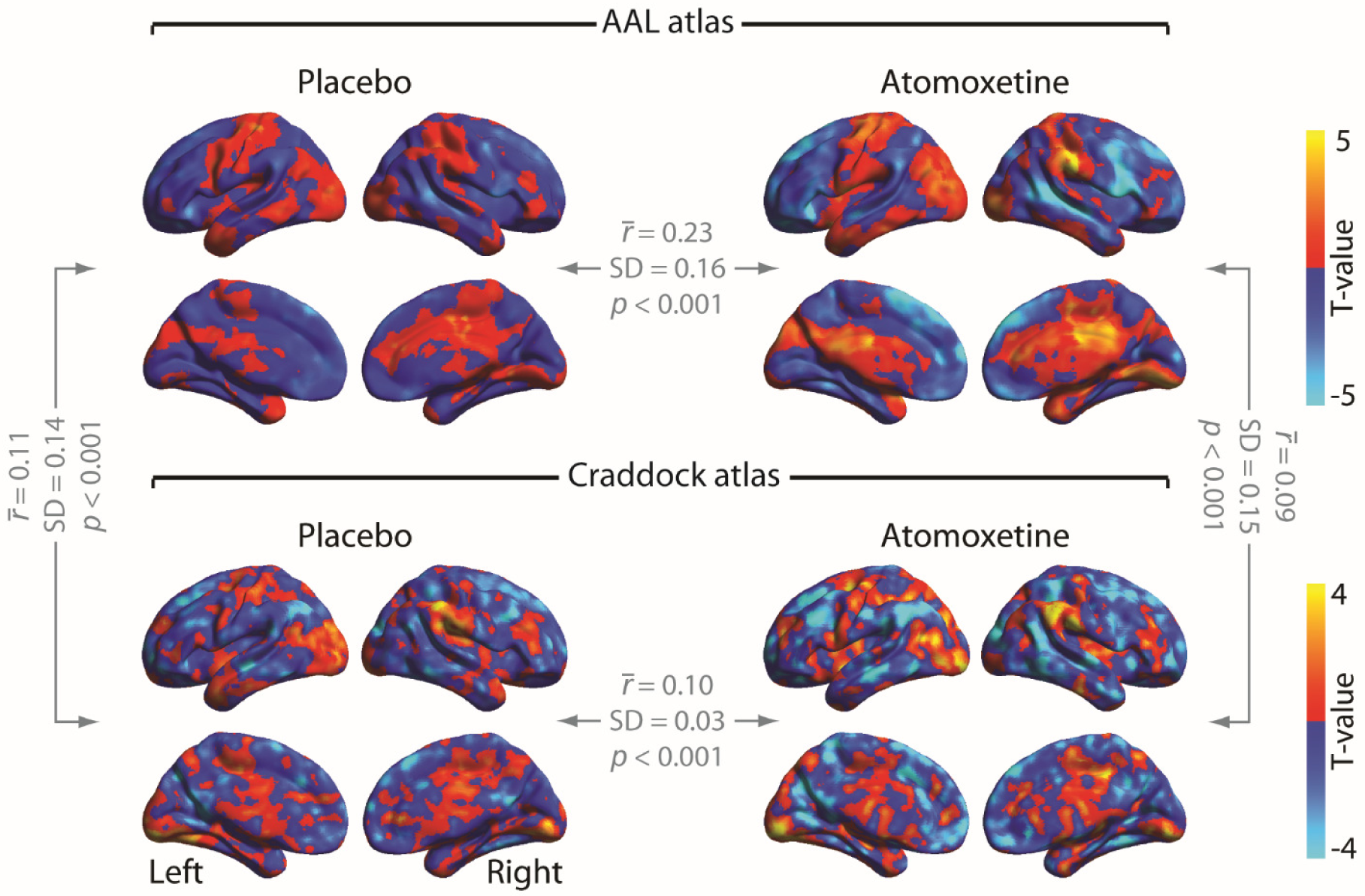
Threshold-free spatial maps of mode 1 for the decomposition atomoxetine > placebo. The 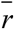 values indicate the average correlation coefficients across participants.

**Figure 8.**
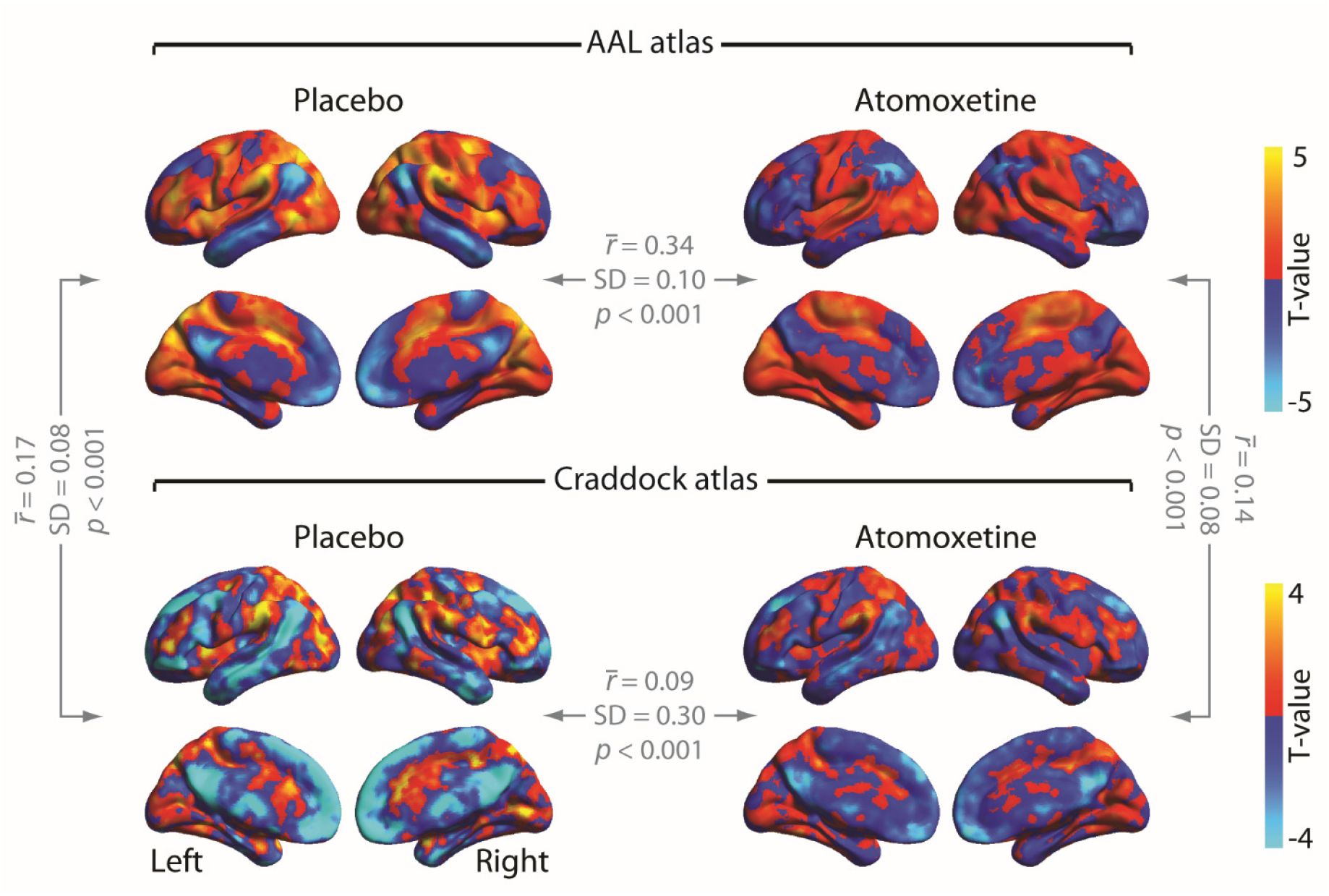
Threshold-free spatial maps of mode 1 for the decomposition placebo > atomoxetine. The 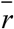 values indicate the average correlation coefficients across participants.

### Control 4: Artifacts in global signal do not account for main results

Recent findings have suggested that the global MRI signal may contain artifacts that are related to various non-neural sources, and these artifacts are not effectively removed by standard preprocessing techniques (Power et al., 2017). Such artifacts may have caused spurious differences between conditions in the structure of inter-regional covariance. We therefore applied global signal (the mean of all regional time series) regression to the regional BOLD time series prior to computing covariance matrices, and repeated our key spatial mode decomposition analyses.

For the decomposition atomoxetine > placebo, the percentage variance explained of mode 1 differed between conditions and in the expected direction (AAL: *t*(23) = 4.45, *p* < 0.001, ROC index = 0.64, *t*(23) = 6.88, *p* < 0.001; Craddock: *t*(23) = 4.55, *p* < 0.001, ROC index = 0.69, *t*(23) = 7.54, *p* < 0.001). For the decomposition placebo > atomoxetine the percentage variance explained of mode 1 also differed between conditions and in the expected direction (AAL: *t*(23) = −5.15, *p* < 0.001, ROC index = 0.63, *t*(23) = 8.97, *p* < 0.001; Craddock: *t*(23) = −6.23, *p* < 0.001, ROC index = 0.63 *t*(23) = 7.06, *p* < 0.001). Moreover, the spatial structure of the modes that included global signal regression was similar to that of the modes that did not include global signal regression, as indicated by significant correlations between mode weights (all r values > 0.42, all *p* values < 0.001). Thus, our findings were unlikely to be driven by spurious differences between conditions relating to artifacts in the global signal.

### Control 5: Differences in peripheral physiology do not account for main results

Because atomoxetine significantly increased both heart rate and breath rate (atomoxetine vs placebo: heart rate: *t*(23) = 3.24, *p* = 0.004; breath rate: *t*(23) = 3.02, *p* = 0.006), it is possible that the RETROICOR denoising procedure operated differently in the atomoxetine and placebo conditions, thereby conceivably introducing spurious changes in the structure of inter-regional covariance. We therefore repeated the spatial mode decomposition analyses on data to which no RETROICOR had been applied. For both atlases and for both decomposition directions, all between-condition comparisons of variance explained by the modes were significant and in the expected direction (AAL, atomoxetine > placebo: p < 0.001; ROC index: 0.62, p < 0.001; Craddock, atomoxetine > placebo: p < 0.001; ROC index: 0.63, p < 0.001; AAL, placebo > atomoxetine: p < 0.001; ROC index: 0.62, p < 0.001; Craddock, placebo > atomoxetine: p < 0.001; ROC index: 0.63, p < 0.001). Moreover, to examine if the modes that resulted from decomposition of non-RETROICOR-corrected data were similar in spatial structure to the modes that resulted from decomposition of RETROICOR-corrected data, we correlated the mode weights between the RETROICOR-corrected and non-corrected modes. All correlations were significant, (all r values > 0.47, all *p* values < 0.001), thus ruling out the possibility that the modes reflected between-condition differences in peripheral physiology.

### Control 6: Eigenvalue decomposition identifies similar networks as ICA

When applied to our dataset, ICA identified components, shown in Figure 9, that corresponded well with so-called ‘resting-state networks’ previously obtained from ICA of fMRI data (Smith et al., 2009). We verified that the linear decomposition approach used here identified similar spatial patterns. To this end, we selected voxel-level mode maps based on maximal spatial correlation with the 10 intrinsic connectivity networks reported by Smith et al. (2009). For the placebo condition, the average correlation coefficient was 0.41 (SD 0.12, min 0.16, max 0.56), and for the atomoxetine condition the average correlation coefficient was 0.40 (SD 0.12, min 0.15, max 0.53). Similar results were obtained with the Craddock atlas.

**Figure 9.**
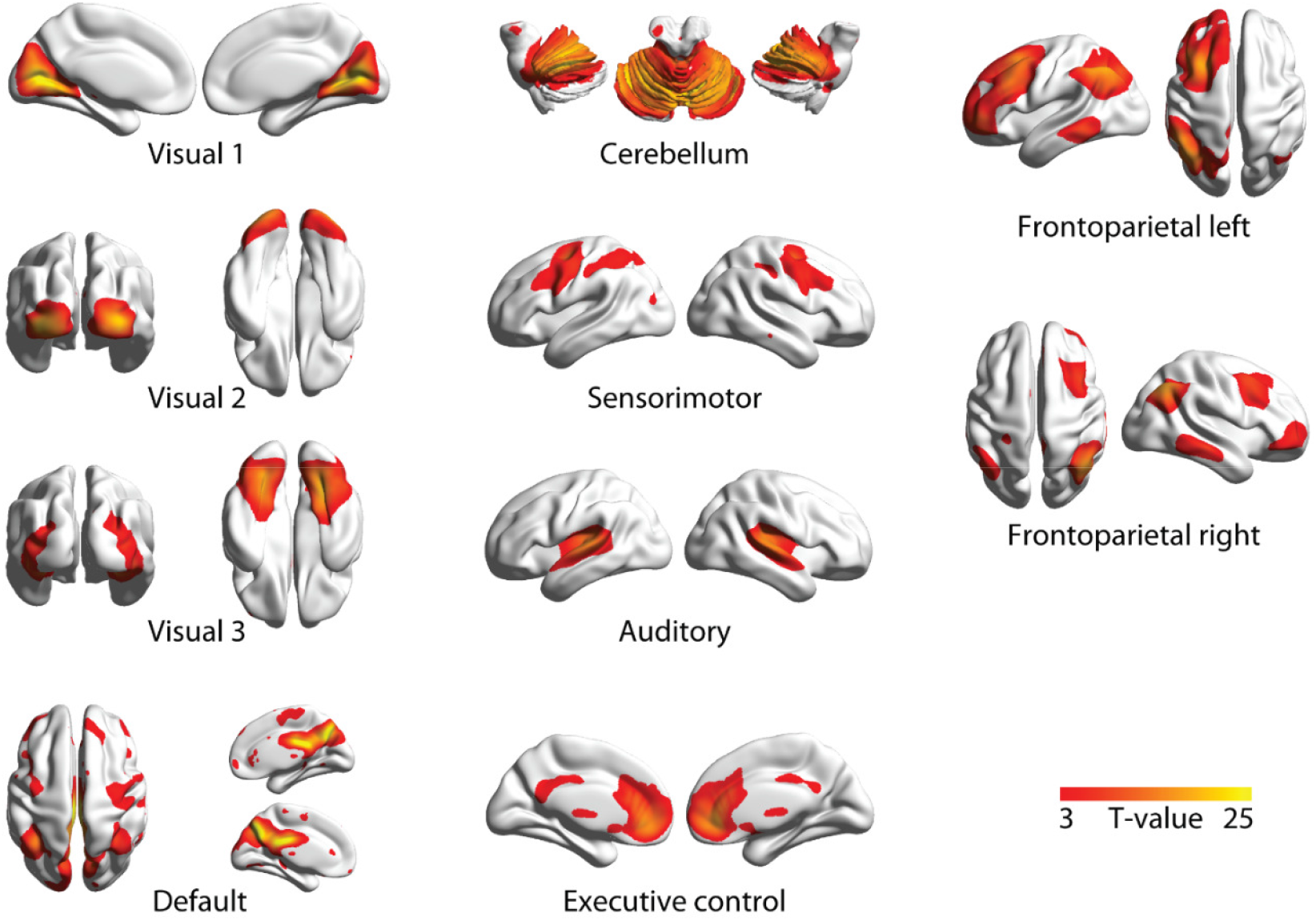
Spatial maps of the independent components that were selected based on spatial correlation with the 10 canonical resting-state networks presented by Smith et al. (2009). Spatial maps were visualized with BrainNet Viewer (Xia et al., 2013).

## DISCUSSION

Catecholamines are important regulators of behavior and have profound effects on physiological brain states, and play a key role in mental disorders (Montague et al., 2004; Aston-Jones and Cohen, 2005; Robbins and Arnsten, 2009; McGinley et al., 2015). A substantial body of work has characterized the catecholaminergic modulation of single neuron activity (Berridge and Waterhouse, 2003; Winterer and Weinberger, 2004) or micro-circuit operations (Marder, 2012; Polack et al., 2013). Fewer studies have assessed catecholaminergic modulation of large-scale brain network dynamics. Pharmacological fMRI studies in monkeys and humans have shown that catecholamines alter the strength of correlations between distant brain regions (Hermans et al., 2011; Guedj et al., 2016; van den Brink et al., 2016; Warren et al., 2016; Hernaus et al., 2017). While ‘resting-state’ studies have reported catecholamine-induced decreases in correlation strength (van den Brink et al., 2016; Guedj et al., 2017), task-based studies have reported increases (Warren et al., 2016; Hernaus et al., 2017), or the converse for noradrenergic antagonism (Hermans et al., 2011). Critically, the brain-wide distribution of these modulatory effects has thus far remained unknown.

Here, we imaged the brain-wide distribution of catecholamine-induced changes in intrinsic correlations across the human brain and related the resulting spatial patterns of brain dynamics to the brain-wide distribution of specific catecholamine receptors. We thus applied an analysis approach tailored to delineate spatial patterns of both drug-induced increases and decreases in correlation strength (Figure 1) to ‘resting-state’ fMRI data from a placebo-controlled atomoxetine intervention. This uncovered two distinct, and widely distributed, sets of brain regions (Figures 2,3), each of which showed a distinct spatial correspondence to the brain-wide distribution of catecholamine receptor genes, but not acetylcholine or NMDA receptor genes (Figure 4). Our results establish that the impact of catecholamines on brain network dynamics exhibits remarkable spatial specificity. Our results bridge between the endogenous modulation of large-scale brain network dynamics and the low-level properties of the underlying neurotransmitter systems.

The catecholaminergic system is equipped with a large variety of receptor types, which are non-uniformly distributed across the cortex (Zilles and Amunts, 2009; Nahimi et al., 2015; Salgado et al., 2016). These receptors have dissociable effects on neural activity (McCormick et al., 1991; Robbins and Arnsten, 2009; Noudoost and Moore, 2011; Salgado et al., 2016). In particular, α_1_ and β receptors have relatively low affinity for NE and are therefore activated only at relatively high synaptic NE levels (e.g., due to stress). These receptors seem to weaken cortical circuit interactions (Ramos and Arnsten, 2007), an interesting observation given that the spatial distribution of these receptors was specifically associated with the spatial mode that captured a catecholamine-induced *suppression* of fMRI signal correlations. By contrast, the spatial mode atomoxetine > placebo (i.e., *enhancement* of correlations) was associated with the expression of D_2_-like receptors, which have been associated with cortical disinhibition (Seamans et al., 2001; Winterer and Weinberger, 2004), and some of which also show particularly high affinity for NE (Arnsten, 2011). Thus, it is possible that inhibition / disinhibition of local populations of neurons cause, by virtue of widespread receptor expression, large-scale decreases / increases in correlation strength respectively. Regardless of the precise mechanistic origin of changes in correlation strength, our findings suggest that the diversity in distribution and function of catecholamine receptors is responsible, at least in part, for the opposite sign modulations of correlations we uncovered here.

This insight is in accordance with the emerging view of the LC-NE system as a more specific regulator of brain-wide neural interactions than traditionally assumed. In addition to the receptor heterogeneity across the brain that we focused on here, recent results indicate that the ascending projections of the LC are more spatially specific than once thought (Chandler and Waterhouse, 2012; Chandler et al., 2014; Schwarz and Luo, 2015; Schwarz et al., 2015; Uematsu et al., 2015; Kebschull et al., 2016; Uematsu et al., 2017). Furthermore, distinct subpopulations of LC neurons mediate opposite behavioral effects (Uematsu et al., 2017), and could thus also affect the underlying neural interactions in dichotomous ways.

In our previous work, we identified atomoxetine-related reductions in signal correlation strength at the whole-brain level (van den Brink et al., 2016). Other fMRI work has revealed similar global changes in the strength of correlations, due to pupil-linked arousal (Eldar et al., 2013; Warren et al., 2016), pharmacological intervention (Hermans et al., 2011; Warren et al., 2016), and concurrent alterations in the topological properties of whole-brain cofluctuations (Shine et al., 2016; Shine et al., 2017). At first glance such unitary modulations of correlations may appear to be at odds with the opposing atomoxetine-related effects in different sets of brain regions that we identified here. However, our analysis approach was specifically tailored to delineate the *predominant* catecholamine-induced changes in fluctuations. Thus, our findings do not rule out the possibility of spatially homogenous modulations of correlations due to catecholamines - they only show that such potential global effects accounted for a smaller proportion of variance than the spatially-specific catecholamine-related changes focused on here. Our current findings should thus be viewed as complementary to previous work, offering a detailed view of the predominant aspects of catecholamine-modulated correlations.

The brain-wide effects of catecholaminergic manipulation observed here stand in striking contrast to the recently reported effects of a cholinergic manipulation (deactivation of the nucleus basalis). The latter attenuates the so-called “global MRI signal” (i.e., averaged across all gray matter voxels) at rest while leaving the structure of specific resting-state networks relatively unaltered (Turchi et al., 2018). Instead, we found that the catecholamine-induced effects are heterogeneous, affecting specific functional networks. Thus, the catecholaminergic and cholinergic systems - despite similarly widespread ascending projections - may have dissociable influences on large-scale brain activity.

Noteworthy is that both spatial modes exhibited a negative correlation between homotopic brain regions. Similar left-right asymmetries in endogenous NE concentration (Oke et al., 1978) and noradrenergic modulations of correlations (Grefkes et al., 2010) have previously been reported. The ‘bilaterally-opponent’ structure we observed (Figures 2b,3b) may have resulted from modulation of inter-hemispheric anatomical connectivity, via direct or indirect pathways, as homotopic brain regions are strongly interconnected (Segraves and Rosenquist, 1982; Lim et al., 2012). In this scenario, catecholamines simply modulated the functional efficacy of the structural connectome. Another (non-mutually exclusive) possibility is that this bilaterally-opponent structure resulted from the spatial structure of the unilateral ascending projections from the left and right LC to the cortex. In addition, atomoxetine shifted correlations from left-lateralized to right-lateralized frontoparietal networks, an observation corroborated by correlation with ICA-derived ‘resting state networks’. Indeed, right-lateralized frontoparietal regions might be particularly susceptible to NE-influences and involved in goal-oriented stimulus processing (Corbetta and Shulman, 2002; Corbetta et al., 2008). It is tempting to speculate (participants were not engaged in a task) that our current results indicate an atomoxetine-related shift towards goal-oriented stimulus processing, a hypothesis that could be tested in future work.

The current study showcases the utility of generalized eigenvalue decomposition for the analysis of resting-state fMRI data. One of its primary advantages over conventional analysis techniques (e.g. dual regression, (Beckmann, 2009)) is that it does not require an *a priori* selection of functional networks, but instead yields the spatial modes that show the strongest drug effects. Thus, it increases the sensitivity to potentially more subtle drug-related changes, as evidenced by the atomoxetine-induced increases in correlated fluctuations that were not identified in our previous study (van den Brink et al., 2016). The approach also has limitations. First, although we demonstrated robustness of results across two particular parcellation schemes, the resulting spatial modes might differ for other parcellation schemes, in particular those of radically different densities. Second, the approach can only be used to compare correlations between two conditions (or groups), limiting its applicability for more complex (e.g., longitudinal) study designs. Third, the approach required focusing on one or a few out of the large number of spatial modes yielded by the decomposition.

An examination of all potential modes of interest revealed that several modes other than the first exhibited statistically significant differences between condition (Figure 5b,d). Each of these modes may have captured meaningful information about atomoxetine-related changes in correlations. We focused on the first mode for a number of reasons. First, orthogonality between the spatial modes that is imposed by the analysis could obscure the interpretation of modes subsequent to the first. Second, for both decomposition directions, the first spatial mode tended to account for more variance in independent data than subsequent modes (Figure 5a-d). We thus used it as a readout of the predominant effect of atomoxetine on correlation strength. Third, for both decomposition directions, the first spatial mode captured the strongest association with the distribution of specific catecholamine receptors (Figure 5e,f).

In sum, we have shown that catecholamines increase and decrease the strength of intrinsic fMRI signal correlations within two distinct sets of distributed brain regions. These spatially-specific and opposite-polarity modulations of ongoing brain dynamics mirror the spatial receptor diversity within the catecholaminergic system. Our results provide a reference for understanding catecholaminergic effects on network interactions during task performance, and important constraints for modeling catecholaminergic effects on the forebrain.

## Acknowledgements

The authors would like to thank Serge Rombouts for his helpful advice, Christopher Warren, Peter Murphy, and Daphne Tona for their help with data collection, Henk van Steenbergen and Janna Marie Bas-Hoogendam for their help with preprocessing the data, and the members of the Donnerlab for valuable discussion. This work was supported by a fellowship for postdoctoral researchers funded by the Alexander von Humboldt foundation (to R.L.v.d.B.), a European Research Council Starting Grant (to S.N.), and the following grants from the German Research Foundation (to T.H.D): DO 1240/3-1 and SFB 936/A7.

